# Genetic Basis of Variation in Cocaine and Methamphetamine Consumption in Outbred Populations of *Drosophila melanogaster*

**DOI:** 10.1101/2021.03.01.433403

**Authors:** Brandon M. Baker, Mary Anna Carbone, Wen Huang, Robert R. H. Anholt, Trudy F. C. Mackay

**Affiliations:** Department of Genetics and Biochemistry and Center for Human Genetics, Clemson University, 114 Gregor Mendel Circle, Greenwood, SC 29646; The Center for Integrated Fungal Research and Department of Plant and Microbial Biology, North Carolina State University, Raleigh, NC 27695; Department of Animal Science, Michigan State University, East Lansing, Michigan 48824

**Keywords:** extreme QTL genome wide association mapping, Drosophila Genetic Reference Panel, RNA interference, advanced intercross populations

## Abstract

We used *Drosophila melanogaster* to map the genetic basis of naturally occurring variation in voluntary consumption of cocaine and methamphetamine. We derived an outbred advanced intercross population (AIP) from 37 sequenced inbred wild-derived lines of the *Drosophila melanogaster* Genetic Reference Panel (DGRP), which are maximally genetically divergent, have minimal residual heterozygosity, are not segregating for common inversions, and are not infected with *Wolbachia pipientis*. We assessed consumption of sucrose, methamphetamine-supplemented sucrose and cocaine-supplemented sucrose, and found considerable phenotypic variation for consumption of both drugs, in both sexes. We performed whole genome sequencing and extreme QTL mapping on the top 10% of consumers for each replicate, sex and condition, and an equal number of randomly selected flies. We evaluated changes in allele frequencies among high consumers and control flies and identified 3,033 variants significantly (*P* < 1.9 × 10^-8^) associated with increased consumption, located in or near 1,962 genes. Many of these genes are associated with nervous system development and function, and 77 belong to a known gene-gene interaction subnetwork. We assessed the effects of RNA interference (RNAi) on drug consumption for 22 candidate genes; 17 had a significant effect in at least one sex. We constructed allele-specific AIPs which were homozygous for alternative candidate alleles for 10 SNPs and measured average consumption for each population; nine SNPs had significant effects in at least one sex. The genetic basis of voluntary drug consumption in Drosophila is polygenic and implicates genes with human orthologs and associated variants with sex- and drug-specific effects.

**Significance Statement:** The use of cocaine and methamphetamine presents significant socioeconomic problems. However, identifying the genetic underpinnings that determine susceptibility to substance use is challenging in human populations. The fruit fly, *Drosophila melanogaster,* presents a powerful genetic model since we can control the genetic background and environment, 75% of disease-causing genes in humans have a fly counterpart, and flies - like humans - exhibit adverse effects upon cocaine and methamphetamine exposure. We showed that the genetic architecture underlying variation in voluntary cocaine and methamphetamine consumption differs between sexes and is dominated by variants in genes associated with connectivity and function of the nervous system. Results obtained from the Drosophila gene discovery model can guide studies on substance abuse susceptibility in human populations.

## Introduction

The use of illicit drugs such as cocaine and methamphetamine contributes to greater than 200,000 deaths worldwide, with an annual economic cost of over $740 billion in the U.S. alone (1, 2). The neurobiology and mode of action of cocaine and methamphetamine is well known. Both drugs are psychostimulants that act on the dopaminergic projection from the ventral tegmental area to the nucleus accumbens. Cocaine blocks the reuptake of dopamine by binding to dopamine transporters on presynaptic neurons (3, 4). Methamphetamine increases release of dopamine from presynaptic vesicles by reversing the flow of vesicular monoamine transporters (5).

Cocaine and methamphetamine use lead to suppressed appetite resulting in malnutrition, arousal, hyperactivity and, with chronic use, cardiovascular and respiratory disorders (6–9). Susceptibility to substance use disorders varies among individuals in human populations and is determined by both genetic and environmental factors. However, our knowledge of the genetic basis of susceptibility to psychostimulants is limited (10). Animal model studies have focused primarily on manipulating genes based on *a priori* knowledge, focusing mainly on genes associated with neurotransmission in the mesolimbic reward pathway (11–16). Although monoaminergic systems play a role in the molecular mechanisms of response to psychostimulant intake, drug abuse phenotypes are highly polygenic and encompass diverse physiological manifestations. Human genome wide association (GWA) studies identified genes and allelic variants associated with drug abuse phenotypes (17–19). However, such studies face challenges and limitations due to the difficulty in recruiting subjects because of criminalization of substance abuse, and hence relatively small samples sizes; variation in drug exposure; heterogeneity in genetic backgrounds and environments; and comorbidity with other neuropsychiatric disorders such as alcoholism.

*Drosophila melanogaster* presents a powerful genetic model since the genetic background and environment can be precisely controlled, drug consumption can be measured in large numbers of flies (20), and 75% of disease-causing genes in humans have a fly ortholog (21). Cocaine and methamphetamine bind to the central binding site of the *D. melanogaster* dopamine transporter (dDAT) (22), and flies exhibit many of the effects that are observed in humans upon cocaine and methamphetamine exposure, such as increased arousal, suppressed sleep and decreased food intake (23, 24).

The *D. melanogaster* Genetic Reference Panel (DGRP) is a collection of 205 fully sequenced, inbred lines derived from a natural population (25, 26). A GWA study that measured consumption of cocaine and methamphetamine in a subset of the DGRP lines revealed genetic variation for drug consumption (20). Furthermore, DGRP derived-outbred populations can be constructed to circumvent limitations of the statistical power inherent in the small sample size of the DGRP, as illustrated by a previous study on the genetic basis of ethanol consumption (27).

Here, we utilize an advanced intercross population (AIP) derived from 37 DGRP lines that are maximally homozygous, unrelated, and free of inversions or infection by the symbiont *Wolbachia pipientis*, to dissect the genetic underpinnings for variation in consumption of cocaine and methamphetamine. We report the use of allele-specific AIPs as an experimental strategy that can validate causality of individual allelic variants associated with variation in cocaine or methamphetamine consumption identified in GWA studies.

## Results

### Phenotypic variation in drug consumption

To determine the optimal concentration to assess variation in consumption of cocaine and methamphetamine, we used the Capillary Feeder (CAFE) assay (27–30) (Figure 1). We quantified voluntary consumption of 4% sucrose and 4% sucrose supplemented with 0.05, 0.1, 0.2, 0.3, 1.0 and 3.0 μg/μL cocaine or methamphetamine for three-to five-day-old AIP flies (Dataset S1). Overall consumption decreased with increasing concentrations of both drugs for both sexes, with a concentration of 1.0 μg/μL near the inflection point of the sigmoidal distribution for both methamphetamine and cocaine (Figure S1).

**Figure 1:**
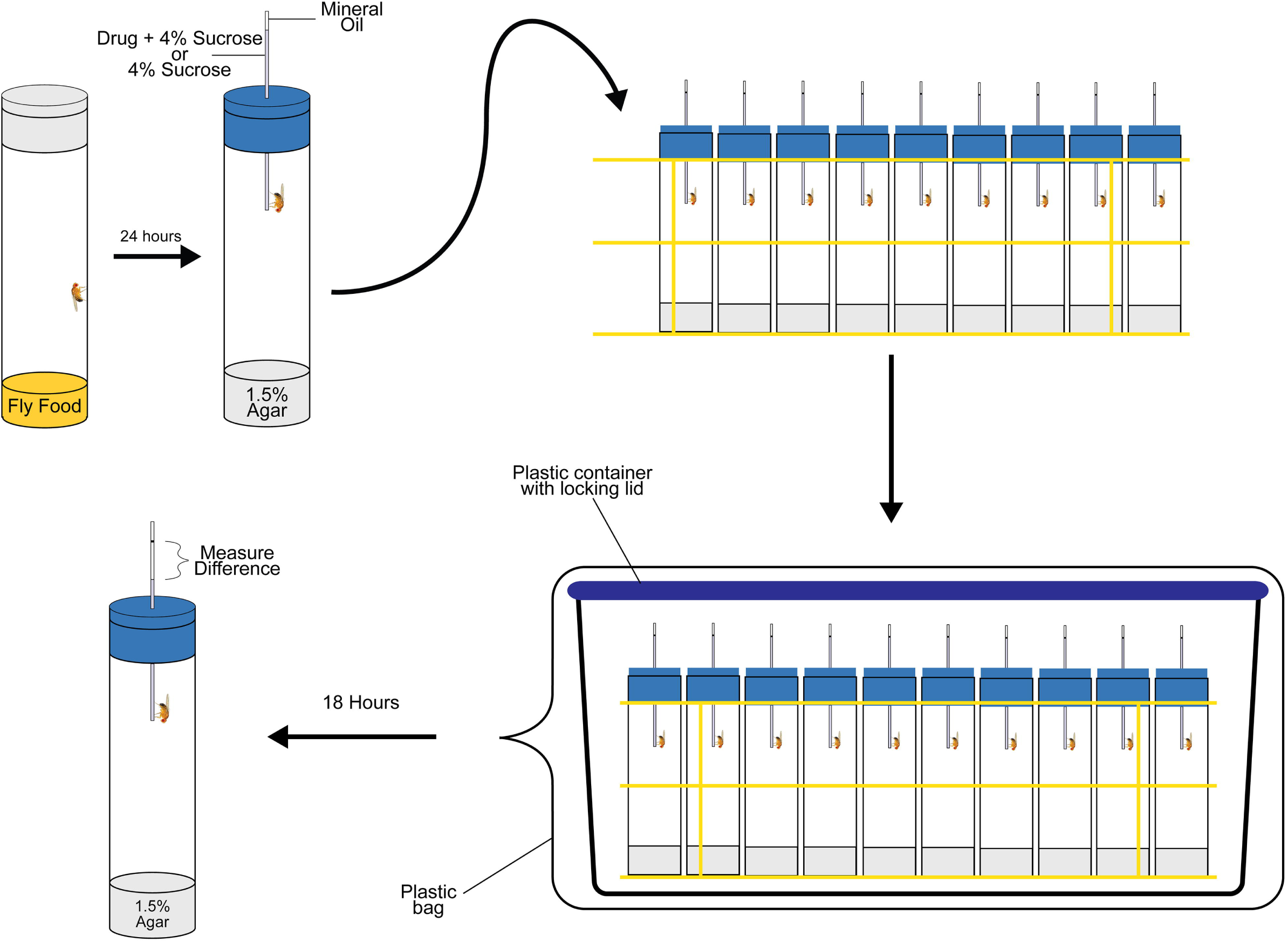
Capillary Feeder (CAFE) assay. Individual flies are collected and placed in vials containing standard fly medium 24 hours prior to beginning the experiment. Flies are then transferred to vials containing 1.5% agar medium, plugged with a foam plug. A capillary filled with the appropriate treatment and a mineral oil cap is then inserted through the foam plug. The initial liquid level is marked on each capillary. Vials are placed in wire racks and the racks are placed inside a plastic container. The plastic container is placed inside a large transparent bag to create a humidified chamber. After 18 hours of feeding, the final liquid levels are marked, and the differences between the initial and final liquid levels are quantified.

Using 1.0 μg/μL cocaine or methamphetamine, we evaluated voluntary consumption of sucrose, sucrose plus cocaine and sucrose plus methamphetamine (hereafter designated the sucrose, cocaine and methamphetamine treatments, respectively) in the AIP. We phenotyped two replicates of 1,500 flies for each treatment and sex, for a total of 18,000 individual flies (Dataset S1). There was substantial variation in consumption of cocaine, methamphetamine, and sucrose in the AIP. The phenotypic distribution for each condition and sex, however, was right skewed with most flies consuming little to no solution, especially for cocaine (Figure S2). This large proportion of low-consuming flies is likely due to a combination of factors: some flies may not learn to drink from the capillaries; flies may have an aversive gustatory response to the drug solutions; and flies may choose not to consume.

### Gustatory aversion

To determine the impact gustatory aversion has on voluntary consumption of cocaine and methamphetamine, we measured proboscis extension responses (PER) (31). Flies extend their probosces in response to a palatable tastant, but not to an aversive tastant. We quantified the PER for naïve AIP males and females in response to sucrose, 1.0 μg/μL cocaine and 1.0 μg/μL methamphetamine (100 flies per sex and treatment) (Dataset S2). We found a significant decrease in the PER to both drugs in both sexes when compared to sucrose (Chi-square *P* < 0.05, Figure S3A). However, the gustatory aversion is not large enough to explain the proportion of flies that did not consume cocaine or methamphetamine in the CAFE assay. Thus, natural variation in gustatory aversion to cocaine and methamphetamine may contribute in part to the observed variation in voluntary consumption but is not likely to be the major contributing factor.

Next, we assessed whether prior consumption of cocaine and methamphetamine affects the propensity for PER. We assessed consumption of 1.0 μg/μL cocaine (240 females, 181 males) and 1.0 μg/μL methamphetamine (238 females, 211 males) for naïve AIP flies, and then measured their PER (Dataset S2). For methamphetamine, in both sexes, there was no significant difference in consumption between flies that extended their probosces (“extenders”) and those that did not (“non-extenders”) (male *P* = 0.28; female *P* = 0.72; Figure S3B). For cocaine, there was no significant difference in consumption between extenders and non-extenders in males (*P* = 0.07; Figure S3B). In females, extenders consumed less cocaine than non-extenders (*P* =0.0051; Figure S3B), indicating consumption does not correlate with propensity for PER.

### Extreme QTL (xQTL) GWA analyses

To explore the genetic underpinnings of drug consumption, we performed whole genome sequencing on the pools of high consumers and random flies to identify differentially segregating alleles. We collected and pooled the top 10% of consumers for each treatment, sex, and replicate (150 flies total per sex/condition), as well as 150 random flies per treatment, sex, and replicate. We compared allele frequencies in the high consuming pools with their respective controls and considered SNPs with a Bonferroni corrected *P*-value of *P* < 1.9 × 10^-8^ significant for each comparison (Dataset S3). In females, we identified 152, 225, and 731 SNPs in or near 128, 169, and 563 genes with significant differences in allele frequency between the high consuming and control pools for the methamphetamine, cocaine and sucrose treatments, respectively (Figure S4). In males, we identified 871, 60, and 999 SNPs in or near 700, 48, and 686 genes with significant differences in allele frequency between the high consuming and control pools for the methamphetamine, cocaine and sucrose treatments, respectively (Figure S4). Across all comparisons, we identified 3,033 SNPs with significant differences in allele frequencies between high consuming and control pools, mapping within 1 kb of 1,962 genes. We observe little overlap at the SNP level and a greater amount of overlap at the gene level between the different sexes and treatments (Figure 2A). The majority of the overlap in genes involved sucrose consumption in either females or males, which may be a result of common genetic determinants of general food intake. Of the 988 genes identified in the drug conditions, 650 (65.6%) possess a putative human ortholog (DIOPT ≥ 3) (Dataset S4).

**Figure 2:**
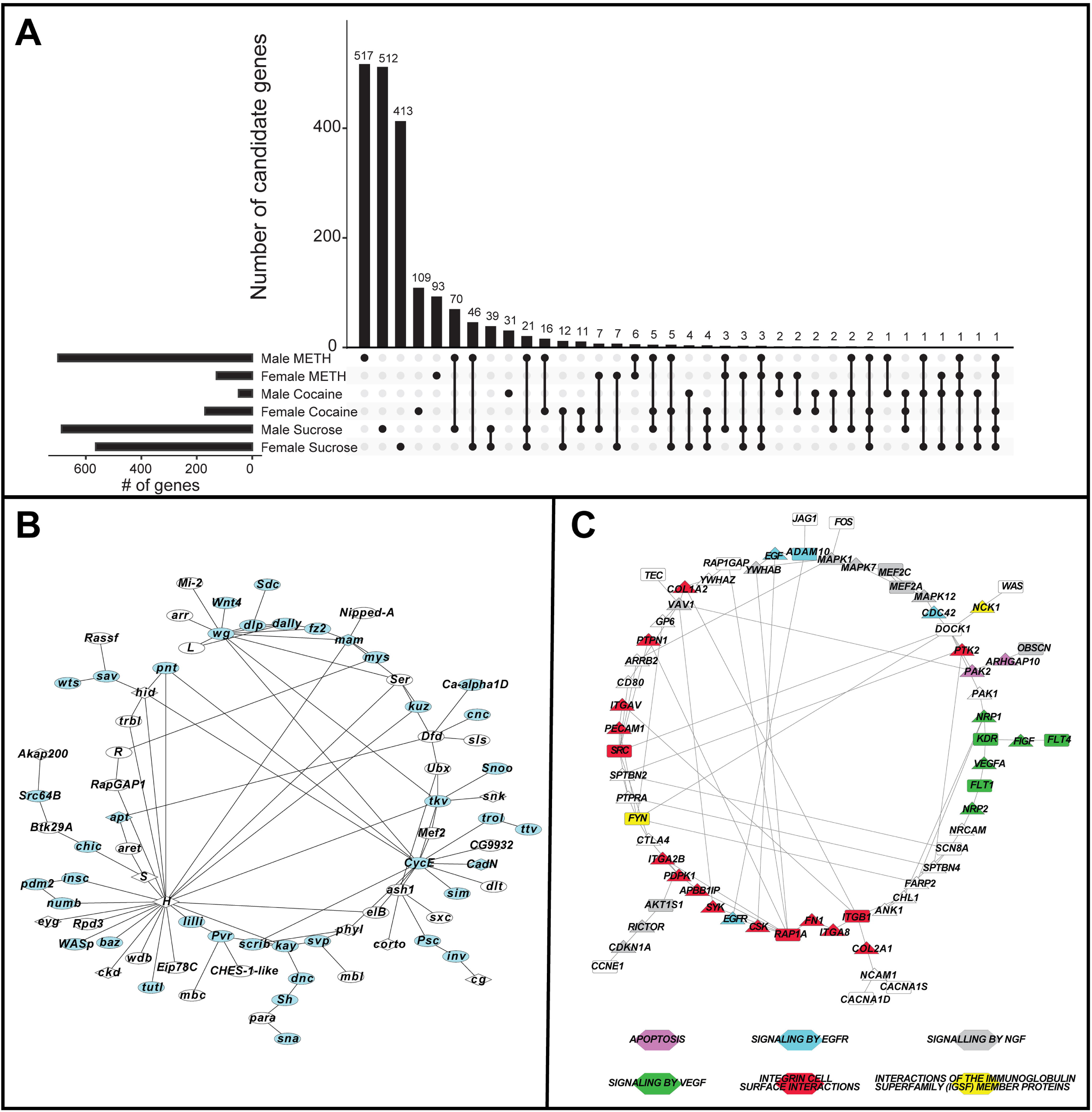
Genes associated with methamphetamine and cocaine consumption. (A) Numbers of genes with allele frequencies that are significantly different between high consuming flies and random control flies. The corresponding number of treatments with which genes are associated is given on the *x*-axis by the dots and interconnecting vertical lines. The horizontal bars indicate the number of significant genes for each treatment. (B) Genetic network without missing genes constructed from a subset of candidate genes identified in the xQTL mapping analysis at a Bonferroni corrected significance level. The network consists of 77 interconnected genes (*P* < 0.001), of which greater than 83% possess a human ortholog (oval nodes; DIOPT ≥ 3). Blue nodes indicate genes associated with nervous system development or function. (C) A genetic interaction network constructed from the human orthologs of genes in (B). Rectangles represent the input human orthologs. Triangles represent computationally recruited genes. Colored nodes indicate the gene is involved in apoptosis (purple), epidermal growth factor receptor (EGFR) signaling (blue), nerve growth factor (NGF) signaling (gray), vascular endothelial growth factor (VEGF) signaling (green), integrin cell surface interactions (red), or interactions of immunoglobin superfamily (IGSF) member proteins (yellow).

We performed gene ontology enrichment analysis using the *PANTHER* Overrepresentation Test (32) and the GO Ontology database (33) for all significant genes in both sexes and both drug treatments, for all significant genes in the cocaine or methamphetamine treatments in both sexes, and for sexes and treatments separately. These analyses reveal enrichment for processes involved in development and morphogenesis, and in particular development and function of the nervous system (Dataset S5).

### Genetic interaction networks

We asked whether genes identified in the xQTL analysis were known to interact in previously curated genetic interaction networks. Without computationally recruiting missing genes, we identified a significant (*P* = 0.001) network with 77 genes, 83.1% of which contain human orthologs (DIOPT ≥ 3) (Figure 2B). Enrichment analysis of this network reveals enrichment of genes associated with elements of nervous system development, including axon extension and guidance, apoptosis; and the Wnt, hippo, Notch, and BMP signaling pathways (Dataset S6).

Using the human orthologs of the genes in the network mentioned above, we constructed a human genetic interaction network by allowing interactions to be mediated by genes not present in our list (Figure 2C). This network is comprised of 66 genes, of which 22 are orthologs of genes in the *Drosophila* genetic interaction network. Multiple biological processes are represented in the network including apoptosis, epidermal growth factor receptor (EGFR) signaling, nerve growth factor (NGF) signaling, vascular endothelial growth factor (VEGF) signaling, integrin cell surface interactions, and interactions of immunoglobin superfamily (IGSF) member proteins.

We also constructed a significant (*P* < 0.001) genetic interaction network in which we integrated 197 genes from a DGRP GWA study on cocaine and methamphetamine consumption (20) with the significant genes from the current study (Figure S5). As expected, the genes in this network are enriched for associations with nervous system development, but there is also enrichment for genes involved in response to cocaine (Dataset S7).

### Functional assessment of candidate genes using RNA interference (RNAi)

We utilized RNA interference (RNAi) to test whether reduced expression of candidate genes identified in the xQTL analysis affects consumption. We selected 22 candidate genes based on the following criteria: they are among the top 10 significant genes (*i.e.,* lowest *P*-values) per treatment or they overlap between at least two treatments; they have a human ortholog with a DIOPT score ≥ 5; there are multiple significant SNPs in the same candidate gene; the allele frequency differences between the control and high consumption pools are in the same direction; the differences in allele frequencies between the control and high consumption pools are consistent across replicates; and an RNAi line with no off-target effects is available in the same genetic background from the Vienna Drosophila Resource Center. For each RNAi line and its co-isogenic control, we measured consumption of cocaine, methamphetamine, or sucrose for 40 males and 40 females for each genotype and treatment tested (Dataset S8). We only tested the conditions for which a SNP was significant for at least one sex in the xQTL analysis. Thus, we tested 18 RNAi lines for methamphetamine consumption, 8 lines for cocaine consumption, and 12 lines for sucrose consumption using the fixed-effects ANOVA model *Y = μ + S + G + S*×*G + ε* and the reduced model *Y = μ + G + ε* for sexes separately, where *Y* is consumption, *μ* is the overall mean, *S* is sex (male or female), *G* is genotype (control or RNAi) and *ε* is the error term.

RNAi targeting of 17 genes had nominally significant *G* and/or *S*×*G* effects in the full model ANOVAs for cocaine and methamphetamine consumption (ANOVA, *P* ≤0.05; Dataset S9). Overall, we validated 77% of candidate genes with differential allele frequencies in the AIP populations that are associated with differences in psychostimulant consumption using RNAi knockdown. A total of 11 of these genes had significant *S*×*G* effects, indicating genetic variation in sexual dimorphism, which can occur if the effect of RNAi on drug consumption is significant in only one sex (sex-specific), if the effects are significant and in the same direction in both sexes but of different magnitudes in males and females, or if there are opposite effects on drug consumption in males and females (sex-antagonistic). Five of the genes (*CG11619*, *qless*, *Pvr*, *RagA-B*, *sim*) had sex-antagonistic effects and two (*CG9348*, *hoe1*) had sex-specific effects on methamphetamine consumption; four additional genes (*arr*, *Cip4*, *Nipped-A*, *nito*) had sex-specific effects on cocaine consumption. All other genes affecting drug consumption were significant in only one sex, even though the *S*×*G* effects were not significant (Figure 3A, 3B). In accordance with the GWA analyses, genetic variation in sexual dimorphism is a hallmark of cocaine and methamphetamine consumption in Drosophila.

**Figure 3:**
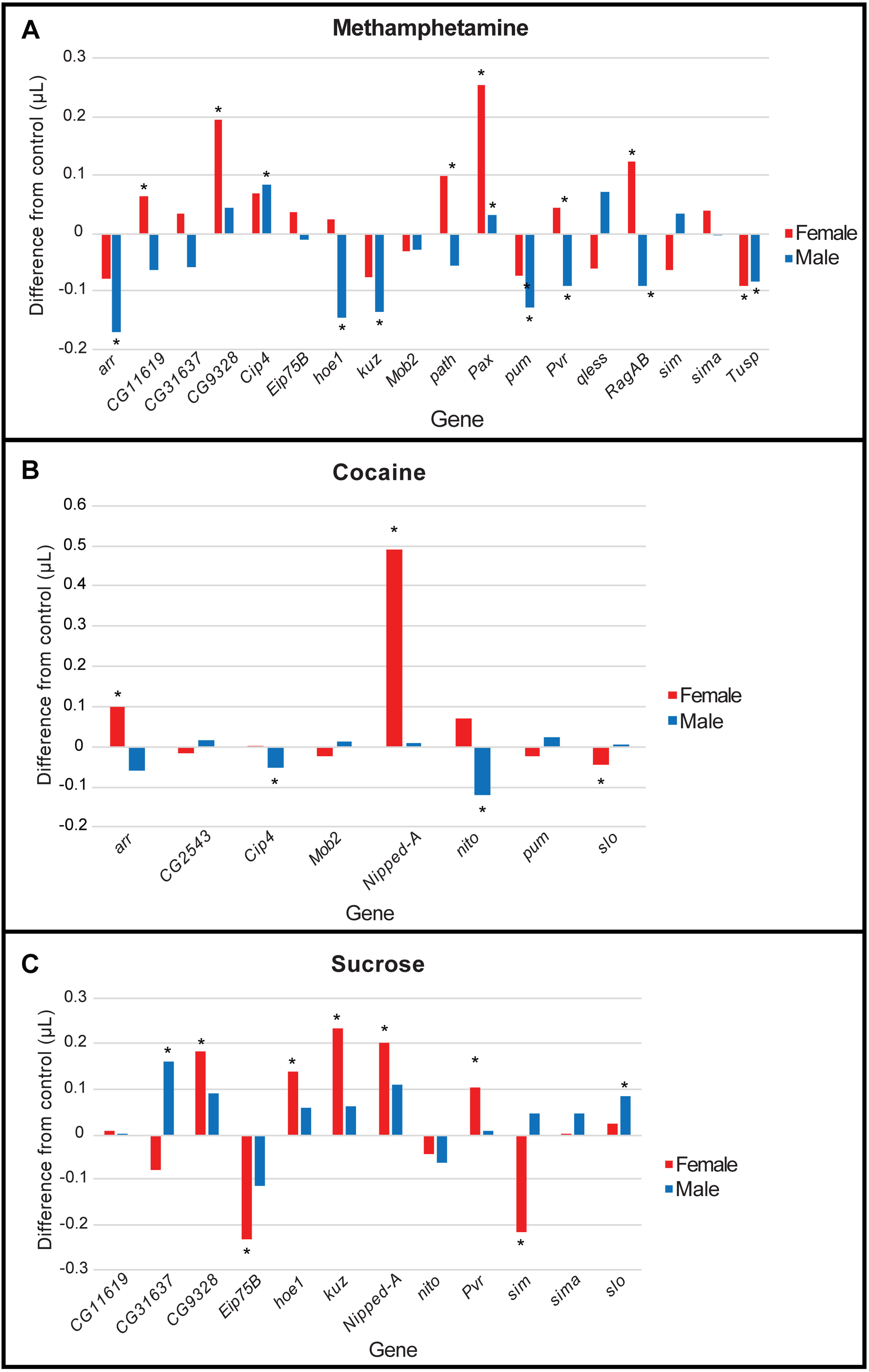
Differences in methamphetamine (A), cocaine (B), and sucrose (C) consumption between RNAi and control genotypes for 22 candidate genes. Data are presented as the difference between the amount consumed in the RNAi line and the amount consumed in the control line. Deviations below zero indicate less consumption in the RNAi line versus the control and deviations above zero indicate greater consumption in the RNAi line. Blue bars denote males and red bars denote females. Bars with an asterisk indicate a significant difference from control (ANOVA, *P* < 0.05).

We tested consumption of both cocaine and methamphetamine for four of the 22 candidate genes (*arr*, *Cip4*, *pum*, and *Mob2*). Cocaine and methamphetamine consumption are significantly different for all four genes (ANOVA treatment (*T*) term, *P* < 0.05; Dataset S9) and for all but *Mob2*, the genotypes respond differently to cocaine and methamphetamine, as indicated by the significant *G*×*T* term. Similarly, we tested nine genes for consumption of methamphetamine and sucrose and three genes for consumption of cocaine and sucrose. Sucrose and drug consumption are significantly different for all of these genes (ANOVA *T* term, *P* < 0.05; Dataset S9). Seven genes showed genotype-specific differences in consumption of drugs and sucrose (significant *G*×*T* or *G*×*T*×*S* terms), while five genes had similar effects of RNAi on drug and sucrose consumption (Dataset S9). Similar to the GWA results, the effects of RNAi of candidate genes on consumption is both shared and distinct between cocaine and methamphetamine, and between drugs and sucrose.

### Functional assessment of candidate SNPs

Whereas RNA interference can corroborate GWA results associated with cocaine and methamphetamine consumption at the level of genes, this method cannot be used to functionally evaluate candidate variants. In addition, SNPs that are in intergenic regions and have large phenotypic effects cannot be validated using publicly available RNAi constructs. To circumvent these limitations, we designed a SNP validation assay by taking advantage of the extensive annotation of variants in the DGRP. We constructed a different set of allele-specific AIPs where alternative SNP alleles were made homozygous in an otherwise randomized genetic background. To independently assess the effects of individual SNPs on cocaine or methamphetamine consumption, we constructed each allele-specific AIP by choosing 10 DGRP lines with the high consumption allele (H) and 10 DGRP lines with the alternative control allele (C) implicated by the xQTL GWA analysis for each SNP of interest. None of the DGRP lines used to construct any of the allele-specific AIPs were the parental lines used to construct the AIP for the xQTL GWA analyses. The 10 H allele DGRP lines and the 10 C allele DGRP lines were separately crossed in a round-robin crossing scheme and maintained for at least 35 generations of random mating in large populations before testing cocaine and methamphetamine consumption in the C and H AIPs.

We constructed allele-specific AIPs for five intergenic SNPs and SNPs located within 1 kb of five annotated genes. We used the following criteria to select intergenic SNPs: they are among the top 20 significant SNPs (*i.e*., lowest *P*-value) per treatment; their differences in allele frequencies between high consuming and control pools are consistent across replicates; they are located in a transcription factor binding site, repeat region, or known regulatory region; they are in a binding site for more than one transcription factor; and there are at least 10 DGRP lines that contain the “H” or “C” allele. We selected genic SNPs that are among the top 10 significant genes per condition based on their *P*-value, or overlap between at least two treatments; they have a human ortholog with a DIOPT score ≥ 5; if there are multiple SNPs, the allele frequency differences between the high consumers and control pools are in the same direction; the allele frequency differences between the high consumers and control pools are consistent across replicates; they have an effect size greater than 2; the gene is in a GO category associated with nervous system development and function; and there are at least 10 DGRP lines that contain the H or C allele. These populations are designated by an arbitrary SNP number (1–5) or gene name followed by a C or H to indicate homozygosity for the H or C allele (*e*.*g*., *qless*H and *qless*C) (Dataset S10).

For each population, we measured consumption of cocaine and/or methamphetamine for 150 males and 150 females (Dataset S11). We only tested populations for consumption of the drug for which the SNP was significant in the original AIP xQTL GWA analysis (Figure S6). Therefore, all populations except for SNP5 were tested for methamphetamine consumption. SNP5 and *DIP-zeta* were tested for cocaine consumption. We tested whether the alternative alleles for each SNP had significant effects on consumption using two-way factorial analyses of variance, where SNP genotype and sex are the two main effects; we also performed reduced analyses separately for males and females. All but one (*path*) of the allele-specific AIPs showed significant differences in consumption between the alternative alleles (Figure 4, Dataset S12). Interestingly, all other SNP effects were different in males and females. The effects of SNP1, SNP5, *sim* and *DIP-zeta* (cocaine consumption) were sex-specific; and the effects of SNP2, SNP4, *Src64B* and *qless* were sex-antagonistic (Figure 4A-C). The population × sex interaction was not significant for SNP3 or *DIP-zeta* (methamphetamine consumption). The H allele had greater consumption than the C allele for SNP1, SNP3, *sim* and *DIP-zeta* (cocaine and methamphetamine consumption) and for one of the two sexes for SNP2, SNP4, *Src64B* and *qless* (Figure 4A-C). The SNP5 H allele had lower consumption than the C allele (Figure 4B-C). Genetic variation in sexual dimorphism for voluntary drug consumption is therefore pervasive not only at the gene level, but also at the level of individual SNPs.

**Figure 4:**
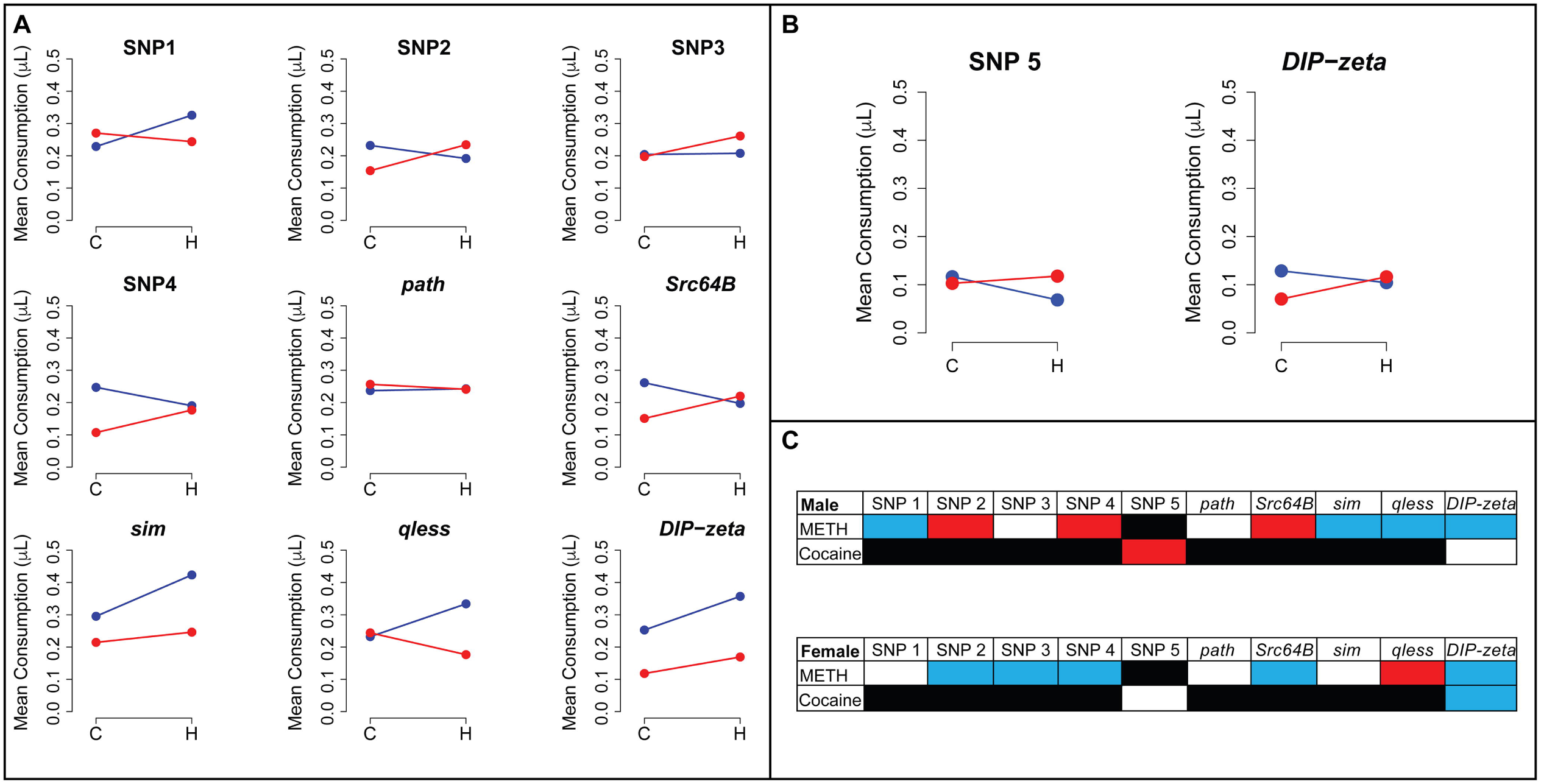
Consumption differences in H versus C allele-specific AIPs. Mean consumption of methamphetamine (A) or cocaine (B) is shown for H and C allele-specific AIP populations. Blue dots represent average consumption for C or H males; red dots represent average consumption for C or H females. (C) Differences in consumption of cocaine or methamphetamine (METH) for males and females in H and C populations for each pair of allele-specific AIPs. Black indicates the drug was not tested for that SNP/gene. Blue indicates significantly greater consumption in the H population versus the C population. Red indicates significantly less consumption in the H population.

## Discussion

Rodent models have long been used to study the physiological, molecular, and behavioral effects of cocaine and methamphetamine, but are not well suited for identifying causal variants in multiple segregating genes that contribute to susceptibility to substance abuse because of limitations on sample sizes, extensive linkage disequilibrium in many mapping populations, and cost considerations. *D. melanogaster* has been used previously to dissect the genetic architecture of ethanol consumption (27), alcohol sensitivity (34), and feeding behavior (35). Here, we used extreme QTL mapping in an advanced intercross population of *D. melanogaster* to identify naturally differentially segregating alleles in individuals with high consumption of sucrose supplemented with cocaine or methamphetamine. We identified significant variants within 1 kb of annotated genes and constructed a genetic network associated with increased consumption of cocaine and methamphetamine. We showed that RNAi of a sample of candidate genes from the GWA analyses affected cocaine and/or methamphetamine consumption for 77% of the genes tested. Our allele-specific AIPs functionally implicated intergenic and intragenic SNPs associated with drug consumption. We found extensive genetic variation in sexual dimorphism underlying the genetic architecture of voluntary cocaine and methamphetamine consumption at the level of GWA associations, RNAi constructs, and allele-specific AIPs, in line with previous studies that documented sexual dimorphism for drug consumption traits in *D. melanogaster* (20, 36), rodents (39, 40) and humans (41).

Previous studies on the effects of cocaine and methamphetamine in Drosophila primarily focused on behavioral effects of single gene mutants (42, 43). Previously, we assessed natural genetic variation in drug consumption traits in a subset of DGRP lines (20). However, the small number of lines in that study only enabled detection of common variants with large effects. Using an AIP and extreme QTL mapping increases statistical power and enabled us to identify 1307 variants in 988 genes that are associated with increased consumption of cocaine or methamphetamine, 209 of which were identified in our previous study, including *arr, Eip75B, pum, Tusp, hoe1, Mob2*, and *kuz* (20). These genes are enriched for biological processes involved in nervous system development, including axon guidance, axon extension, dendrite morphogenesis, and synaptic target recognition.

Genes associated with development of the nervous system have been previously implicated in drug consumption traits in flies (20). Development of the nervous system is recapitulated in the genetic interaction network we constructed consisting of 77 genes (Figure 2B), which shows enrichment for developmental signaling pathways, including Wnt, Notch, and Hippo (Supplementary Dataset 6). In mice, Wnt signaling plays a role in cocaine-induced neuroplasticity (44) and cocaine activates Notch signaling by disrupting the blood-brain barrier (45). Of the 77 genes in this network, 64 contain a human ortholog (DIOPT ≥ 3). We can construct an interaction network consisting of these orthologs and computationally recruited genes (Figure 2C), suggesting evolutionary conservation of fundamental biological processes that underlie variation in susceptibility to psychostimulants.

We selected 22 candidate genes to assess whether RNA interference would affect consumption of cocaine or methamphetamine. We found that 15 (68%) of these genes showed a significant difference in consumption of cocaine or methamphetamine in at least one sex after RNAi knockdown using a weak ubiquitous driver. Failure of candidate genes to show an effect on drug consumption upon RNAi knockdown could be due to functional redundancy, insufficient extent of knockdown (noting that the extent of RNAi knockdown is not linearly related to the phenotypic effect), or the candidate gene may be a false positive.

One limitation of RNAi is that it provides a gene-based rather than a SNP-based validation method and cannot be used to establish causality of intergenic SNPs, some of which have large phenotypic effects. To validate causal SNPs that contribute to variation in consumption of cocaine and methamphetamine, we constructed allele-specific AIPs to isolate alternate alleles of candidate SNPs in randomized genetic backgrounds. This provided an effective method to validate both intragenic and intergenic SNPs. All but one of the AIPs showed significant differences in consumption between the alternative alleles in at least one of the sexes. For most SNPs, the population containing the “high consumer” (“H”) allele consumed more drug than the population with the “control” (“C”) allele in at least one sex. In one case, however, the population containing the H allele consumed less drug than the population with the C allele, which may be attribuDataset to epistasis, which is a dominant feature of complex traits (46, 47).

All four genes for which we have validated allelic variants that are associated with variation in cocaine consumption have human orthologs. The *single minded* (*sim*) gene encodes a helix-loop-helix transcription factor which has been identified as a positive regulator of the midline formation in the embryonic central nervous system (48) and plays a role in axon guidance in the larval brain (49). *DIP-zeta* encodes a Dpr-interacting protein. The Drosophila genome contains 21 members of the *Dpr* gene family and nine paralogs that encode *Dpr* interacting proteins. These proteins contain immunoglobulin domains, and unique paired combinations of DPRs and DIPs function as synaptic partners consolidating neural connectivity (50). *Src64B* specifies a cytoplasmic protein tyrosine kinase, which contributes to development of the mushroom body (51). *qless* encodes a protein involved in the synthesis of the isoprenoid side chain of Coenzyme Q, which is a component of the mitochondrial electron transport chain. Neurons of *qless* mutants undergo mitochondrial stress and caspase-dependent apoptosis (52). These observations support the notion that allelic variants of these genes result in variation in neural connectivity and function, which modulates their propensity to consume psychostimulants.

In conclusion, we developed a method for establishing causality of inter- and intragenic SNPs in outbred populations. Using this approach in combination with RNA interference we showed that the genetic architecture underlying variation in voluntary cocaine and methamphetamine consumption is sexually dimorphic and dominated by genes associated with nervous system development. The use of allele-specific DGRP-derived AIPs can be generally applied to establishing causality of single candidate SNPs associated with variation in complex traits. Based on evolutionary conservation of fundamental cellular pathways, results obtained from the Drosophila gene discovery model can guide studies on substance abuse susceptibility in human populations.

## Materials and Methods

### Drosophila stocks

We created an advanced intercross population (AIP) by crossing 37 inbred, wild-derived *D. melanogaster* lines from the *D. melanogaster* Genetic Reference Panel (DGRP) using a round-robin crossing scheme (27,53,54) followed by over 50 generations of random mating prior to commencement of these studies. The founder lines were minimally related, minimally heterozygous, inversion free and free of *Wolbachia* infection. The AIP was maintained with 400 randomly selected males and 400 randomly selected females each generation. Every generation, 40 males and 40 females were randomly selected from each of 10 bottles and re-distributed at random into 10 bottles to minimize genetic drift. Flies were allowed to lay eggs for 24 hours to minimize crowding; the adults were then discarded.

We obtained *UAS*-RNAi lines from the Vienna Drosophila Resource Center (VDRC) for 22 candidate genes (Supplementary Dataset 13). The *P{KK}* RNAi lines are from the same genetic background and contain an upstream activating sequence (*UAS*) construct at the same locus on the second chromosome (55). The *P{GD}* RNAi lines are from the same genetic background and contain *P*-element based transgenes in a random insertion site (56). None of these RNAi lines had predicted off-target effects. We also obtained the co-isogenic control lines for the KK (*w*^1118^, v600100) and GD (*w*^1118^, v60000) RNAi lines from the VDRC. We generated the *Ubi156-GAL4* driver line in house (53). All flies were maintained on cornmeal-agar-molasses medium at 25°C, 70% humidity and a 12h:12h light/dark cycle.

### Capillary Feeder (CAFE) assay

Individual three-to-five-day old flies were placed on culture medium for 24 hours, and then transferred to culture vials containing 1.5% agar medium. Vials were plugged with blue foam plugs in which a 5 μL capillary tube containing either 4% sucrose or 4% sucrose supplemented with cocaine or methamphetamine was inserted. A drop of mineral oil was added to the top of each capillary tube to impede evaporation. Vials were placed in racks in a humidified chamber to further minimize evaporation. Each humidified chamber contained “blank” vials containing a capillary but no fly to quantify the evaporation rate. Fluid level measurements were recorded immediately after the capillaries were inserted into the vials and after 18 hours of *ad libitum* feeding (Figure 1). The decrease in fluid level was converted to volume consumed following normalization for evaporation.

### Dose response

We performed dose response experiments to determine the appropriate concentration of cocaine and methamphetamine for consumption in the AIP. We assessed consumption of sucrose supplemented with 0.05, 0.1, 0.2, 0.3, 1.0 and 3.0 μg/μL cocaine or methamphetamine for 50 males and 50 females. Mean consumption for each concentration was calculated and plotted with an S-curve super-imposed on the means. For each sex and condition, the inflection point was close to a concentration of 1.0 μg/μL. We also performed dose response experiments for the GD and KK RNAi control lines to determine the optimal concentration of cocaine and methamphetamine for RNAi validation. We quantified consumption of sucrose supplemented with 0.25, 0.5, 0.75, and 1.0 μg/μL cocaine or methamphetamine for 30 males and 30 females(Supplementary Dataset 14). Few flies from these genotypes consumed cocaine or methamphetamine at concentrations of 0.75 and 1.0 μg/μL; therefore, we selected 0.5 μg/μL cocaine and methamphetamine for testing the RNAi lines.

### Drug consumption in the AIP

We quantified consumption of sucrose, sucrose supplemented with 1.0 μg/μL cocaine, or sucrose supplemented with 1.0 μg/μL methamphetamine for males and females separately. Each assay day, the top 10% of consumers for each condition and an equal number of random, unexposed flies were separately collected into pools and flash frozen for DNA sequencing. In total, we quantified consumption for 1,500 males and 1,500 females with two replicates per condition for sucrose, cocaine, and methamphetamine consumption (18,000 flies total).

### Proboscis Extension Response (PER)

Three-to-five-day old AIP flies were collected and placed on culture medium for 24 hours. Flies were then transferred to vials containing non-nutritive 1.5% agar for 24 hours, after which they were anesthetized using CO_2_, and mounted on a glass slide with nail polish with the ventral side facing upwards. After a recovery period of 3-6 hours at 25°C, the flies were tested for their proboscis extension response (PER) (31). Flies were satiated with water then presented with tastants for two to three seconds, with five to ten second intervals between tastants. The order of presentation was: water, sucrose, drug, water, sucrose. Individual flies were only tested with either 1.0 μg/μL cocaine or 1.0 μg/μL methamphetamine. The PER was calculated as the percent of flies that extended their proboscis in response to the tastant. Only flies that responded to sucrose initially were recorded. Flies were tested until 100 flies were scored per sex and treatment. Differences in PER between sucrose and cocaine or methamphetamine were assessed using a Chi-squared test.

PER was also analyzed for individual three-to-five-day old males and females following consumption of cocaine or methamphetamine for 18 hours using the CAFE assay. Flies that did not survive or did not respond to sucrose on the initial presentation were not scored. Differences in consumption between extenders and non-extenders were assessed using a two-tailed *t*-test.

### DNA sequencing

Pools of 150 flies per sample were homogenized using a TissueLyser (Qiagen, Inc., Germantown, MD, USA). Genomic DNA was extracted with the Gentra Puregene Tissue kit (Qiagen Sciences, MD, USA) then fragmented to 350-400 bp using a Covaris S220 Sonicator (Covaris, Inc, Woburn, MA, USA). 100 ng of fragmented DNA were used to produce barcoded DNA libraries using NEXTflex^TM^ ChIP-seq barcodes (Bioo Scientific, Inc., Austin, TX, USA). Libraries were quantified using Quant-IT dsDNA HS Kits (ThermoFisher, Waltham, MA) on a SpectraMax Plate reader and their sizes (bp) were determined using the 2100 Bioanalyzer (Agilent Technologies, Inc., Santa Clara, CA, USA). All libraries were diluted to 10 nM and all libraries for each replicate were pooled together, denatured, and diluted to 16 pM. Pools were clustered on an Illumina cBot (Illumina, Inc., San Diego, CA, USA) and sequenced on eight lanes (four lanes per pool) on an Illumina HiSeq2500 sequencer (Illumina, Inc., San Diego, CA, USA) using 125 bp paired-end sequencing. Sequence reads were aligned to the *D. melanogaster* reference genome using Burrows-Wheeler Aligner (BWA, version 0.6.2) (56). Low quality bases at the end were trimmed with the “-q 13” option in BWA. The alignments were locally realigned, marked for PCR duplicates using GATK (version 2.4) (57) and Picard tools (version 1.89) before recalibrating base qualities with GATK. Bases passing a series of quality filters (47) were piled up to obtain counts of alleles at polymorphic sites where the parental lines segregate.

### Extreme QTL GWA analyses

We performed ‘extreme QTL mapping’ (xQTL) (58) to identify variants for which there are consistent differences in allele frequencies between randomly selected (C) and high consumption (H) pools, averaged over both replicates, for sexes separately. We used a Z test where the null hypothesis is that the allele frequencies between the C and H pools are equal and the test statistic compares the difference in the estimated allele frequencies in the C and H pools, taking into account the number of chromosomes in each pool and the sequencing depths in the C and H pools. The test statistic is 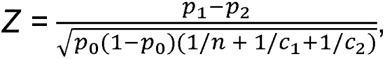 where *p_1_* and *p_2_* are the allele frequencies in the two pools respectively, *p_0_* is the average allele frequency of *p_1_* and *p_2_*, *n* is the number of flies in each pool (*n* = 150), and *c_1_* and *c_2_* are the sequence coverages in the two pools. Under the null hypothesis of no difference between *p_1_* and *p_2_*, *Z* is distributed as standard normal. Evidence for joint association from the two replicate populations was obtained by calculating a combined *χ*^2^ statistic, weighted by sequence coverage, and obtaining *P*-values from the *χ*^2^ distribution (59). We tested 2,568,908 SNPs and used a Bonferroni-corrected significance threshold of 1.95 x 10^-8^ to determine significance of individual variants, separately, for the sexes and three treatments.

### Enriched genetic interaction networks

We annotated candidate genes identified in the xQTL analyses using complete genetic interaction networks from FlyBase (release r5.57) which were curated based on the literature. The genes are nodes in the network and the interactions are edges between the nodes. We mapped significant candidate genes from the xQTL analyses onto this graphical representation of genetic networks and extracted sub-networks involving the candidate genes, with no missing nodes. We tested whether the maximum sub-network is significantly greater than expected by chance using a permutation procedure (34,60,61). We also constructed a genetic interaction network using the candidate genes identified in the xQTL analysis and the candidate genes identified in (20) using the same procedure described above. A genetic interaction network for human orthologs of the candidate genes in the xQTL analyses was constructed using R-Spider (62). All networks were visualized using Cytoscape 3.8.0.

### Gene Ontology enrichment

Gene ontology (GO) enrichment analyses were performed using the *PANTHER* Overrepresentation Test (32) and the GO Ontology database (33).

### Human ortholog identification

Human orthologs were obtained using the DRSC Integrative Ortholog Prediction Tool (63, 64).

### RNA interference (RNAi)

We crossed four or five homozygous *Ubi156-GAL4* driver males to either 4-7 virgin *UAS-*RNAi females or 4-7 virgin females from the appropriate control strain. 40 males and 40 females per genotype and treatment were collected and aged until they were 3-5 days old. Flies were then separated individually into vials containing culture medium for 24 hours before being transferred to vials containing 1.5% agar medium. Consumption of 4% sucrose, 0.5 μg/μL cocaine plus 4% sucrose, or 0.5 μg/μL methamphetamine plus 4% sucrose was then tested, males and females separately, using the CAFE assay. Only the treatments that were statistically significant in the AIP were tested. Consumption differences between RNAi lines and their corresponding control were assessed separately for each treatment using the fixed-effects ANOVA model *Y = μ + S + G + S*×*G + ε* and the reduced models *Y = μ + G + ε* for sexes separately, where *Y* is consumption, *μ* is the overall mean, *S* is sex (male/female), *G* is genotype (control/RNAi) and *ε* is the error term.

For SNPs/genes where both cocaine and methamphetamine consumption were quantified, we assessed differences in consumption of cocaine and methamphetamine using the fixed-effects ANOVA model Y = *μ + S + G + T + S*×*G + S*×*T + G*×*T + S*×*G*×*T + ε* where *S* and *G* are sex and genotype as mentioned above, and *T* is treatment (cocaine or methamphetamine).

### Allele-specific AIPs and drug consumption

We constructed 10 pairs of allele-specific AIPs: one AIP for the high-consumer (H) allele or the alternate/control (C) allele for each of 10 candidate SNPs or haplotypes. For each allele-specific AIP we chose 10 independent DGRP lines containing the candidate allele that were not used as parental lines for the original AIP. We used the same round-robin crossing scheme described above and maintained all AIPs with 400 randomly selected males and 400 randomly selected females exactly as described above. These lines were maintained for over 35 generations prior to quantification of consumption phenotypes. We tested 150 three-to five-day old flies per sex and condition for consumption of the drug for which the SNP was significant in the original xQTL analysis. To assess differences in consumption between the alleles at each tested locus, we used the two-way factorial fixed effects ANOVA model *Y = μ + S + G + S*×*G + ε* and the reduced model *Y = μ + G + ε* for sexes separately, where *Y* is consumption, *μ* is the overall mean, *S* is sex (male/female), *G* is genotype (C versus H) and *ε* is the error term.

## Acknowledgements and funding sources

Research funded by National Institute on Drug Abuse (U01 DA041613 to TFCM and RRHA)

---

**Figure S1: Dose responses for cocaine and methamphetamine consumption in the AIP.** To determine the optimally discriminating concentrations of cocaine and methamphetamine, we tested consumption of 0.05, 0.1, 0.2, 0.3, 1.0 and 3.0 μg/μL cocaine and 0.05, 0.1, 0.2, 0.3, 1.0 and 3.0 μg/μL methamphetamine for 50 males and 50 females from the AIP. The *y*-axes represent the average consumption for all flies for each concentration. An S curve was superimposed to determine the approximate inflection point. For all four sex/treatment combinations the inflection point falls closest to a concentration of 1.0 μg/μL.

**Figure S2: Distribution of consumption in the AIP.** We tested the consumption of two replicates of 1500 males and 1500 females for cocaine, methamphetamine, and sucrose. The distribution of consumption for all treatments in both sexes is right skewed.

**Figure S3: Proboscis Extension Response (PER).** (A) The PER was tested for cocaine and methamphetamine in 100 AIP males and 100 AIP females. For cocaine, 88% of females and 85% of males extended their proboscis. For both sexes, 91% of flies extended their proboscis for methamphetamine. There is a significant reduction in PER for both sexes and drugs (Chi-square test, *P* < 0.05) compared to sucrose. (B) We tested consumption and PER of cocaine or methamphetamine in males and females from the AIP to determine if the propensity for PER correlates with the previous amount of drug consumed.

**Figure S4: Extreme QTL (xQTL) mapping analysis.** We performed tests for significant differences in allele frequencies between pools of high consuming and random control flies for 2,568,908 SNPs, separately for each treatment and sex. The Manhattan plots show the -log_10_*P* transformed *P*-values for all SNPs. The *x*-axes show the chromosome locations for each SNP. The dashed lines represent the Bonferroni-corrected significance level (P = 1.95 × 10^-8^). Dark red and dark blue dots represent SNPs with a significant difference in allele frequency between control and high consuming pools in females and males, respectively.

**Figure S5: Genetic interaction network for significant genes for high consumption and preference for cocaine or methamphetamine.** We compiled the significant genes from the current study with those from Ref. 20 and constructed a genetic interaction network that consists of 197 interconnected genes (*P* < 0.001) using Cytoscape 3.8.0. Blue nodes indicate genes with a human ortholog (DIOPT ≥ 3).

**Figure S6: Allele-specific AIP testing treatments.** Treatments tested for allele-specific AIPs. Green indicates the drug was tested for that SNP/gene. Black indicates the drug was not tested.

**Dataset S1: Raw consumption data in the AIP.** (A) Dose response raw consumption values. (B) Raw consumption for the xQTL analyses. NA indicates a consumption value outlier that may have been influenced by excessive evaporation.

**Dataset S2: Raw PER data in the AIP.** (A) PER for males and females. (B) PER in males and females after consumption of cocaine or methamphetamine.

**Dataset S3: Extreme QTL (xQTL) GWA analysis in the AIP.** Significant SNPs (following Bonferroni correction for multiple tests) and their allele frequency differences between control and high pools are given for each condition, sex, and replicate. Annotated genes are given for SNPs within 1 kb of the gene body.

**Dataset S4: Human orthologs for candidate genes from the xQTL analyses.**

**Dataset S5: Gene Ontology (GO) enrichment analyses for candidate genes from the xQTL analyses.** (A) All candidate genes from methamphetamine and cocaine treatments. (B) Methamphetamine consumption candidate genes. (C) Cocaine consumption candidate genes. (D) Female methamphetamine consumption candidate genes. (E) Male methamphetamine consumption candidate genes. (F) Female cocaine consumption candidate genes. (G) Male cocaine consumption candidate genes.

**Dataset S6: Gene Ontology (GO) enrichment analysis for 77 candidate genes from the xQTL GWA analyses that belong to a significant (*P* < 0.001) genetic interaction network.**

**Dataset S7: Gene Ontology (GO) enrichment analysis for 197 genes overlapping between the xQTL and DGRP (Ref 20) GWA analyses and that belong to a significant (*P* < 0.001) genetic interaction network.**

**Dataset S8: Raw consumption values for RNAi and control lines.** Group indicates the control line and RNAi lines that were analyzed together. NA indicates an outlier with excessive evaporation.

**Dataset S9: Full and reduced model ANOVAs for candidate genes evaluated for consumption using RNAi.** Significance codes: ***; *P* < 0.001; **; *P* <0.01; *; *P* <0.05. (A) Full model pooled across sexes, methamphetamine. (B) Full model pooled across sexes, cocaine. (C) Full model pooled across sexes, sucrose. (D) Reduced model by sex, methamphetamine. (E) Reduced model by sex, cocaine. (F) Reduced model by sex, sucrose. (G) Full model pooled across methamphetamine and cocaine and sexes. (H) Full model pooled across methamphetamine and sucrose and sexes. (I) Full model pooled across cocaine and sucrose and sexes.

**Dataset S10: Naming scheme for allele-specific AIPs.** SNP locations are from FlyBase Release 5.57.

**Dataset S11: Allele-specific AIP raw consumption data.**

**Dataset S12: Full and reduced model ANOVAs for allele specific AIPs.** Significance codes: ***; *P* < 0.001; **; *P* <0.01; *; *P* <0.05. (A) Full model pooled across sexes. (B) Reduced model by sex.

**Dataset S13: RNAi lines.**

**Dataset S14: Raw consumption values for the dose responses of RNAi control lines.**

