## Supplementary figures and images for "Genetic Basis of Variation in Cocaine and Methamphetamine Consumption in Outbred Populations of *Drosophila melanogaster*"

### Figure S1

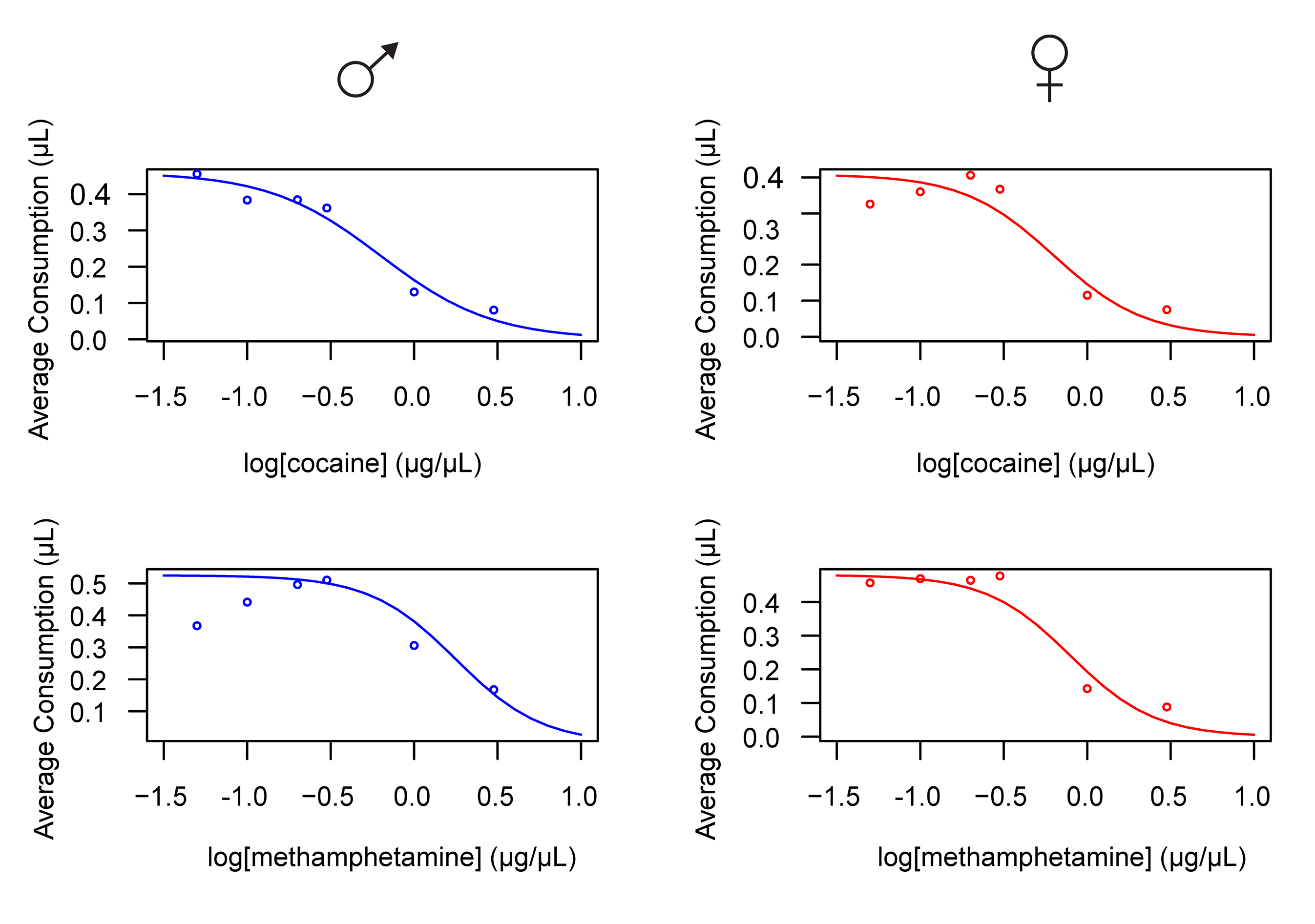

### Figure S2

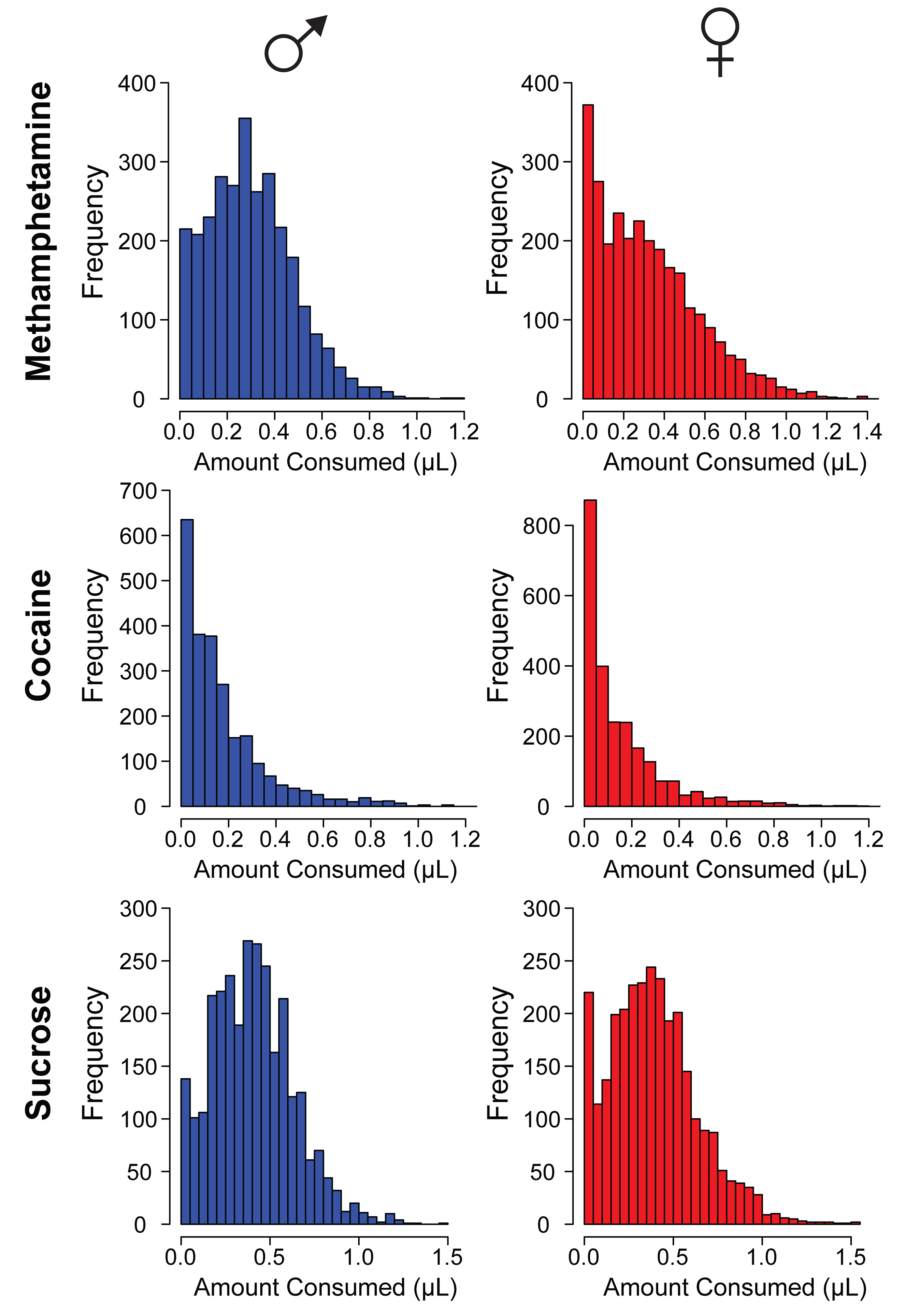

### Figure S3

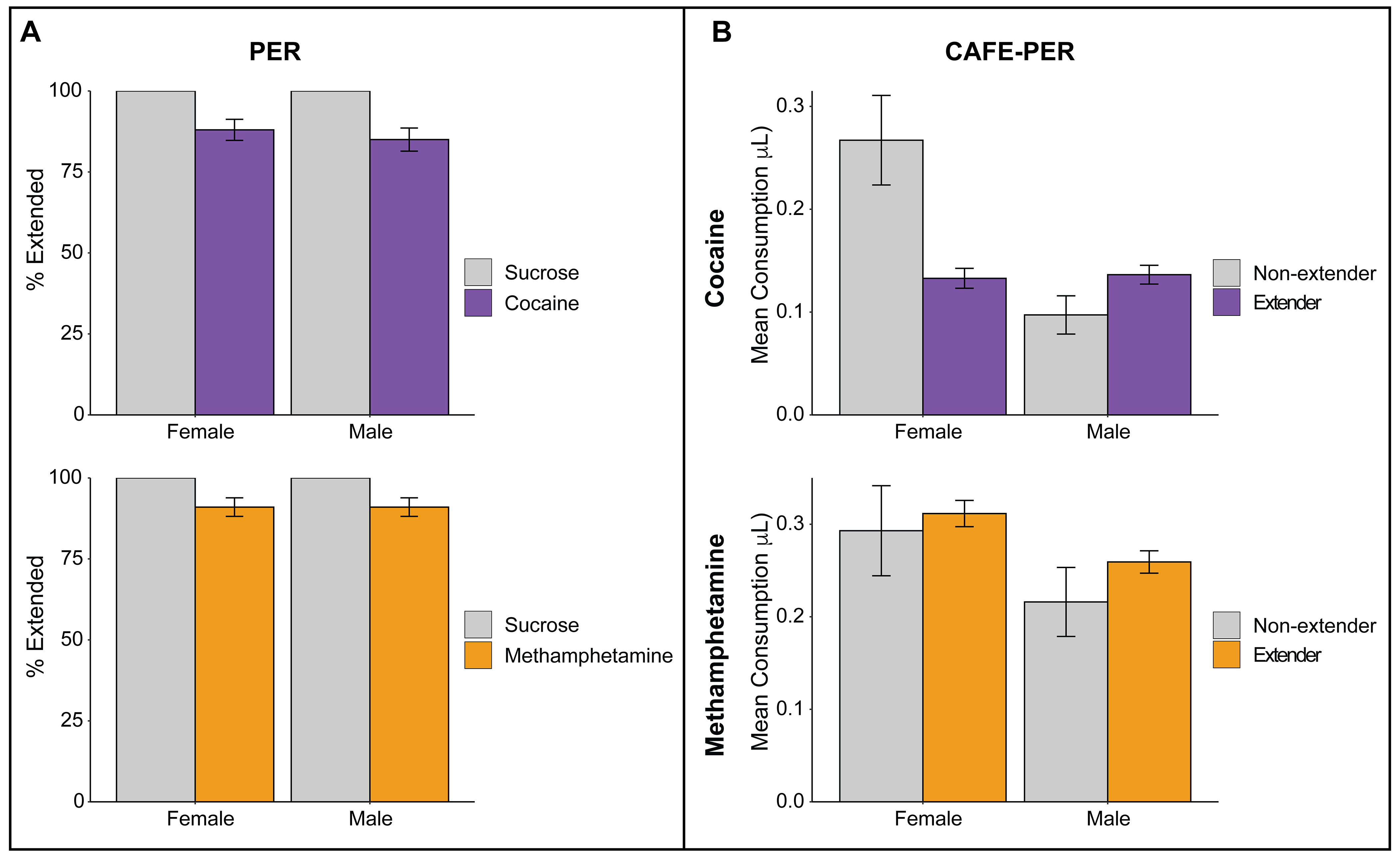

### Figure S4

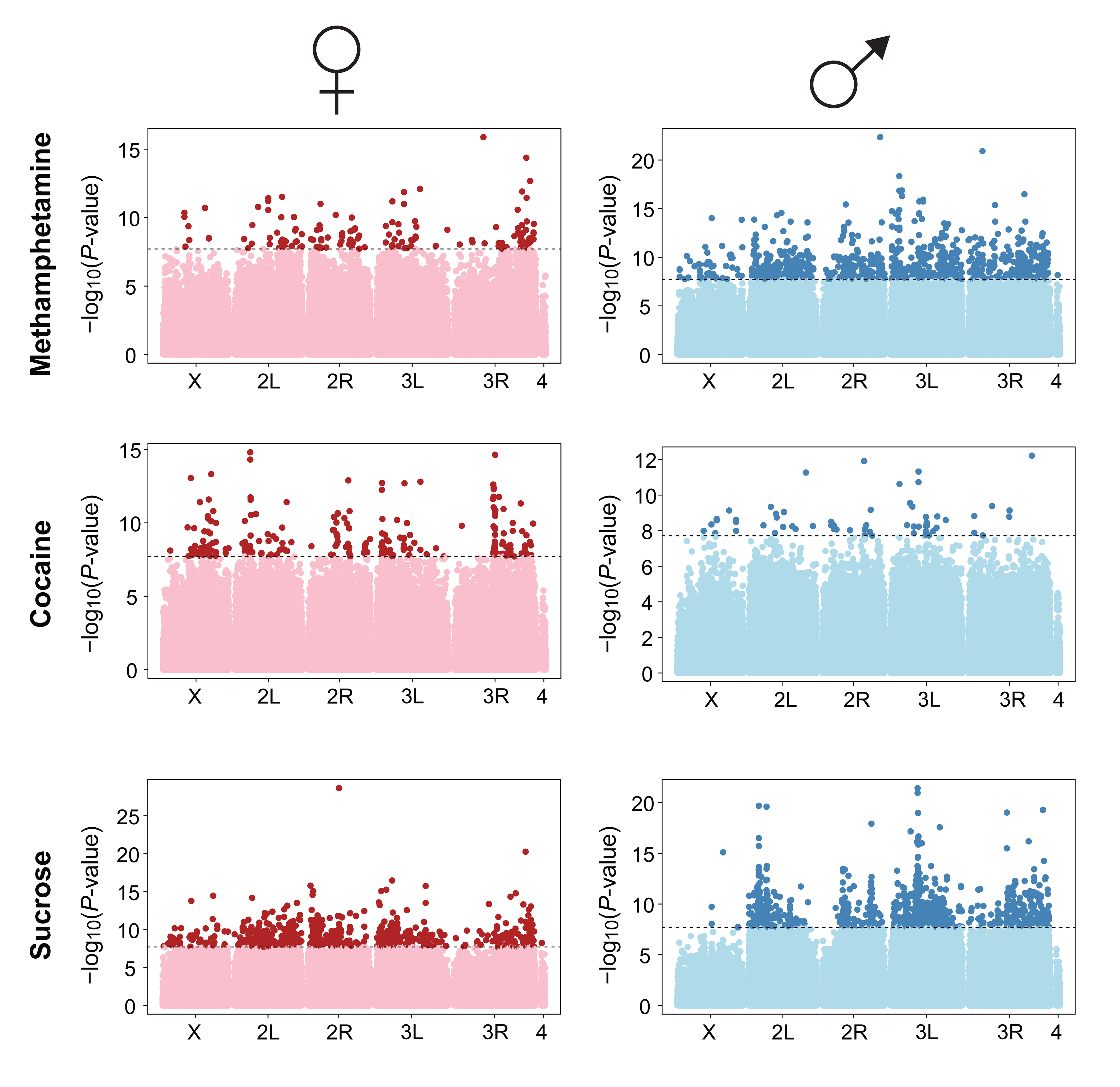

### Figure S5

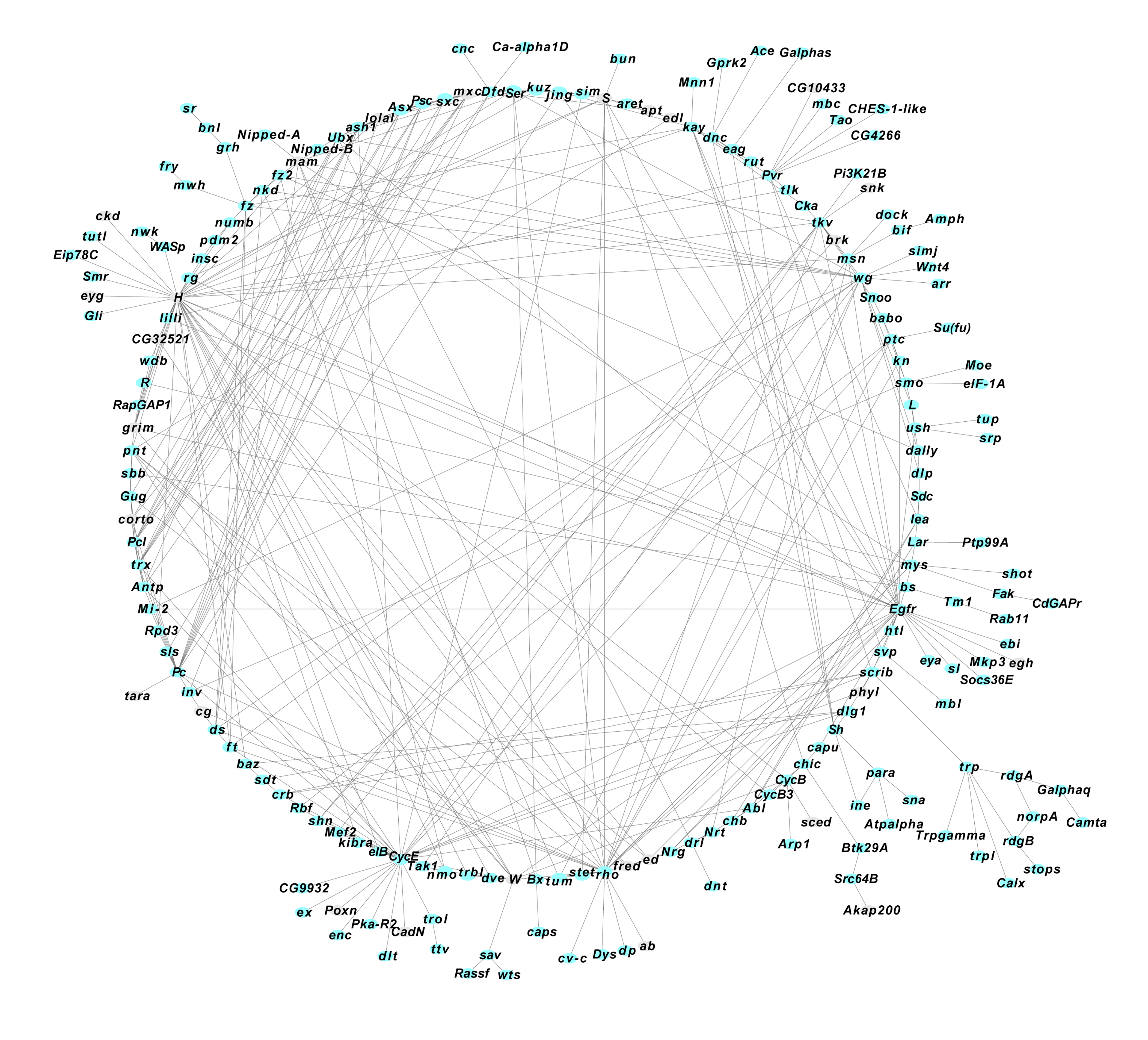

### Figure S6

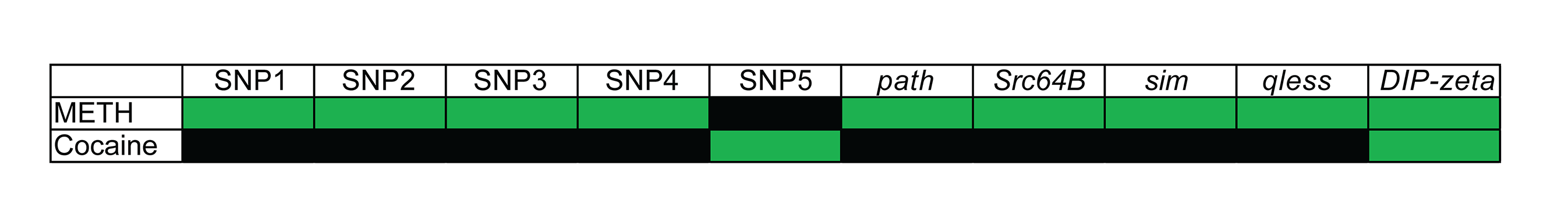
